# Quantitative evaluation of dogs as sentinels for Rift Valley fever virus circulation in Madagascar

**DOI:** 10.1101/2025.05.16.654436

**Authors:** Herilantonirina Solotiana Ramaroson, Andres Garchitorena, Vincent Lacoste, Soa Fy Andriamandimby, Matthieu Schoenhals, Jonathan Bastard, Katerina Albrechtova, Laure Chevalier, Domoina Rakotomanana, Patrick de Valois Rasamoel, Modestine Raliniaina, Heritiana Fanomezantsoa Andriamahefa, Mamitiana Aimé Andriamananjara, Lova Tsikiniaina Rasoloharimanana, Solohery Lalaina Razafimahatratra, Claude Arsène Ratsimbasoa, Benoit Durand, Chevalier Véronique

## Abstract

Sentinel animals may play a key role in surveillance of zoonotic arbovirus circulation, particularly in developing countries with low levels of investment in health. This study aimed to assess the relevance of using dogs as sentinel animals for Rift Valley fever virus (RVFV) surveillance in Madagascar.

Serological surveys were conducted on 513 dogs and 135 cattle in the Ifanadiana district. In addition, 486 human dry blood samples available from the same area were used. Serostatus against RVFV was determined using a competitive Enzyme-Linked Immunosorbent Assay (cELISA) for dog and cattle samples, and a Multiplex Bead Assay (MBA) for human samples. Serocatalytic models fitted to age-stratified serological data were developed to estimate the RVF force of infection (FOI) under several hypotheses, ranging from no relationship to proportional FOIs between humans, cattle, and dogs.

Antibodies to RVFV were detected in 23 of 513 dogs (4.5%; 95% confidence interval (95% CI): [2.9 - 6.7]), in 86 of 486 humans (17.7%; 95% CI: [14.4 – 21.4]), and in 33 of 135 cattle (24.4%; 95% CI: [17.5 - 32.6]). Based on the deviance information criterion, the best supported model indicated that FOI in humans and cattle was proportional to FOI in dogs. Proportionality parameters were estimated at 2.6 (95% credible interval (95% CrI): [1.4 - 5.1]) for humans and 3.5 (95% CrI: [1.3 - 6.4]) for cattle.

Our findings suggest that sampling dogs could be used to identify RVFV circulation in endemic areas, and infer the exposure of humans and cattle in these areas in Madagascar. This original result paves the way for an innovative surveillance method for RVFV in Madagascar and other endemic countries, but also for other arboviruses such as West Nile virus.

**Author Summary:** Rift Valley fever (RVF) is a viral zoonosis transmitted by mosquitoes and through contact with the blood, tissues or body fluids of infected animal. The disease has caused at least three major epidemics in Madagascar to date. RVF mainly affects ruminants through abortion and high mortality rate in young animals, and a range of influenza-like to more severe syndromes that can lead to death in some cases in humans. Dogs that interact with humans and are exposed to mosquito bites in the same environment may not develop clinical signs or sufficient viremia to infect vectors, but they could still develop an immune response when exposed to RVF virus (RVFV). This study aimed to assess the relevance of using dogs as sentinel animals for RVFV circulation on the island. Serocatalytic models were developed using serological data to represent and quantify the relationships between the force of infection (FOI) of RVFV exerted on dogs, humans and cattle. We found a proportional relationship between the FOI exerted on dogs and that exerted on humans and cattle. These findings support the relevance of using dogs as sentinel animals for monitoring the circulation of RVFV and arboviruses in general.

## Introduction

Mosquito-borne viruses have a major impact on public health on a global scale. They are mainly of African origin and some have spread worldwide due to dynamic changes in vector distribution, migratory movements of susceptible hosts and international trade [1]. The most recent arbovirus outbreak in Madagascar occurred in 2021 in the Mananjary district, in the southeastern part of the island, where RVFV was confirmed (Type M immunoglobulin positive) in 16 of 40 suspected cattle, and 12 of 103 suspected humans [2].

RVF is a zoonotic disease caused by an arbovirus belonging to the *Phlebovirus* genus of the *Phenuiviridae* family, within the *Bunyavirales* order [3]. RVF virus (RVFV) was first detected in the Rift Valley in Kenya, in 1930 [4]. Since then, RVFV has been reported in most countries of Africa, the Arabic Peninsula and the Indian Ocean. Outbreaks are characterized by high mortality rates in young ruminants and abortions in adults [4,5]. In humans, symptoms range from self-limiting febrile illness to hepatic and haemorrhagic syndromes, which can lead to death [6].

RVFV was first isolated in Madagascar from mosquitoes in 1979, without any previous record of human or animal cases. Major outbreaks (epidemics and epizootics) occurred in 1990-91 on the east coast and in the central highlands, in 2008-09 on the north and south coasts and in the central highlands, and in 2021 in the southeastern part of the country [2,7–9]. Genetic analysis of viral genomes showed that outbreaks were mainly associated with the reintroduction of the virus in the country from mainland East Africa [10]. However, despite long inter-epizootic periods, persistent low-level virus circulation is suspected to maintain sufficient herd immunity to limit the occurrence of outbreaks [11,12]. Indeed, the enzootic circulation of RVFV in Madagascar could be maintained by ruminant trade in affected areas, and would be supported by simultaneous vectorial and direct transmission, whose intensity could vary between eco-climatic regions [11,13–15]. Entomological studies have shown that 24 mosquito species present on the island were potential vectors of the virus [16,17]. A recent study showed that 10 mosquito species were involved in the last RVFV epidemic/epizootic in the Mananjary district, including 4 species that had never been associated with RVFV transmission in Madagascar before [2,17].

Madagascar is one of the poorest countries in the world, with a poverty rate of 75% of the population in 2022 (below 2 USD per day) according to the World Bank [18]. Resources for human and animal health are limited, making the population particularly vulnerable to epidemics [19]. The island’s high biodiversity is increasingly threatened by human activities such as deforestation and bushfires for agricultural purposes, and natural disasters like cyclones. These factors increase the risk of zoonotic disease emergence. It is thus crucial to identify areas where public health actions need to be prioritized.

For arbovirus serosurveillance, mosquito blood meal screening is a potential tool. However, it requires advanced labs and is limited by the <24h windows post-feeding [20]. Serological surveys are often difficult to carry out in human populations for both ethical and logistical reasons. The use of sentinel animals can be a cost-effective alternative for monitoring zoonotic risks to human health [21]. An ideal sentinel animal should be susceptible to infection, survive it, and develop a sufficient and detectable immune response. Moreover, it should not be able to transmit infection to handlers, nor develop a level of viremia that could infect vectors [22]. According to Bowser et al., domestic dogs (*Canis lupus familiaris*) have these characteristics for many infectious agents of public health importance [23], including arboviruses. Several studies have shown dog exposure to flaviviruses, such as West Nile virus (WNV) [24,25], Dengue virus (DENV) [26], and Japanese Encephalitis virus (JEV) [27]. Two studies have reported RVFV seropositivity in dogs [28,29]. Although these results demonstrate that dogs are exposed to arboviruses in the natural environments they share with humans, none of these studies have quantitatively analyzed the relationship between the exposure of dogs and humans in a given area.

In Madagascar, dogs are present in most communities. Apart from their role as pets, they can serve as guardians and/or be used for hunting. Up to 90% of middle-class households own at least one dog in urban areas (such as Antananarivo) [30], while the average number of dogs per household is higher in rural areas (e.g. 1.55 in Ranomafana and 3.23 in Andasibe) [31]. The vast majority of rural dogs roam freely, and some may interact extensively with the wild environment [30–32]. They also share the same environment with humans and are therefore exposed to the same pathogens [33].

The aim of this study was to examine the potential role of dogs as sentinels for RVFV in Madagascar by quantitatively analyzing the relationship between the exposure of dogs, cattle and humans to RVFV. To achieve this goal, we used available human sera and collected blood samples from dog and cattle populations in the same area of south-east Madagascar. These samples were serologically analyzed for RVFV antibodies using a commercial competition enzyme-linked immunosorbent assay (cELISA) for dogs and cattle, and a multiplex powered by Luminex technology for humans. We then used serocatalytic models to test for a relationship between the Force of Infection (FOI) of dogs and that of humans and cattle, and to quantitatively evaluate the parameters of this relationship.

## Materials and methods

### Study site

The study was conducted in the district of Ifanadiana, a rural district in the Vatovavy region in south-eastern Madagascar (Fig 1). The region, district, commune and *fokontany* are Madagascar’s first, second, third and fourth (smallest) administrative units, respectively. The Ifanadiana district covers an area of 3,970 km^2^, comprises 15 communes and 195 *fokontany* and is characterized by deep valleys and a mountainous landscape with an altitude gradient of 100 to 1,000 meters from East to West [34,35].

**Fig 1.**
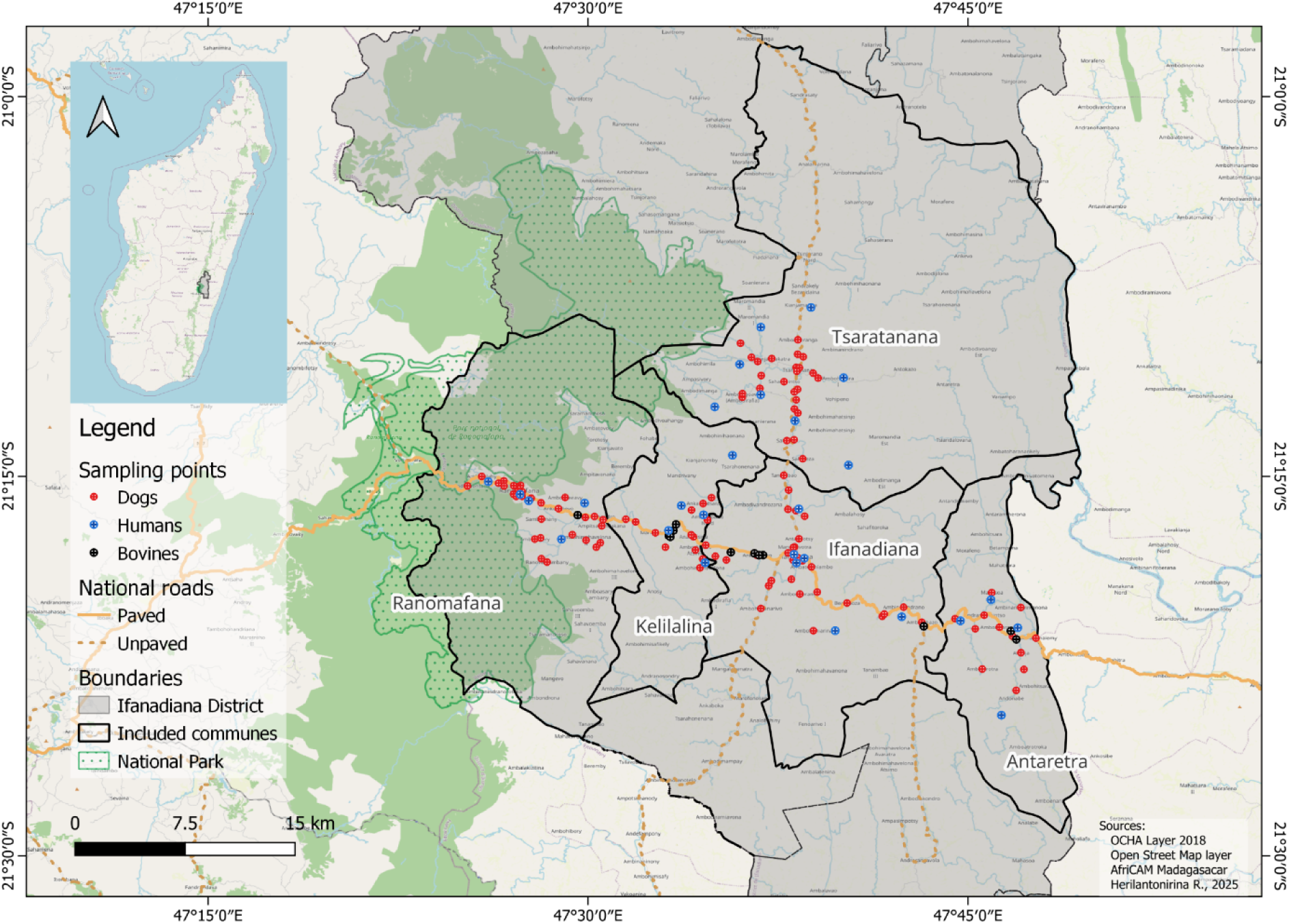
Location of human, dog and cattle sampling points in the Ifanadiana district, Madagascar.

The landscape features a protected area of dense rainforest in the western part (Ranomafana national park), alongside large agricultural fields and steppe areas. Recorded total rainfall in the district is 1,900 mm on average per year, and temperatures vary between 15°C and 32°C depending on altitude and season [36]. With an estimated population of 183,000 inhabitants in 2020, rice production is the main agricultural activity [37].

Three host species were considered for this study: humans, dogs and cattle. Although the last reported RVF outbreak occurred in the Mananjary district, also located in the Vatovavy region [2], we chose to carry out our study in 5 communes of the neighboring district of Ifanadiana located 115 km from Mananjary. This choice was firstly driven by the availability of human samples and associated survey data from an existing longitudinal cohort study (“IHOPE”) [38] described below, and secondly, by the possibility of sampling dogs and cattle with the help of local veterinarians and local community health workers in the remote areas. The selection of these communes finally took into account their accessibility. These communes were Ranomafana, Kelilalina, Ifanadiana and Antaretra, crossed from west to east by National Road 25, and Tsaratanana which is crossed from south to north by the main unpaved road of the district (Fig 1).

### Ethical statements

The IHOPE cohort study was approved by the Madagascar National Ethics Committee - *Comité d’Ethique de la Recherche Biomédicale auprès du Ministère de la Santé* - CERBM) and the Harvard Medical School Institutional Review Board (IRB, N° IRB16-0347). An amendment was obtained for changes in the 2021 surveys and collection of dried blood spot samples (CERBM - N°024 MSANP/SG/AGMED/CNPV/CERBM; 26 February 2021). Use of these samples for the current study was authorized in 2023 (N° 53 MSANP/SG/AMM/CERBM). The dog study was approved by the *Comité d’Ethique National Animal* (CENA - N° 01-23/CENA; 30 May 2023) and has obtained research authorization from the *Ministère de l’Environnement et du Développement Durable (MEDD -* N° 237/23/MEDD/SG/DGGE/DAPRNE/SCBE.Re; 11 July 2023).

Regarding to cattle, the study was approved by and performed with the support of the *Direction des Services Vétérinaires* (DSV)*, Ministère de l’Agriculture et de l’Elevage*.

### Serological data

To ensure that the sampled humans, dogs and cattle had been exposed to the RVFV during equivalent periods of time, we targeted children and animals born after 2010, and at least of 3 months old to avoid potential interference with colostral immunity (S1 Table).

### Dog data

A cross-sectional survey was carried out in dogs from July to September 2023. To determine the sample size, we assumed that seroprevalence is lower in dogs than in cattle, in which it had been estimated at 19.3% in animals aged 1 to 12 years [11]. Based on an expected prevalence of 10%, we used Cochran’s formula to set the number of animals to be sampled at 35 dogs per year of birth (for a relative precision of 100% and assuming perfect sensitivity and specificity of the serological test).

We carried out door-to-door visits and identified households owning at least one dog with the support of community health workers and administrative officers (*Fokontany* chiefs). Accessible *Fokontany* included communes within a maximum walking distance of 5km from the main road were surveyed. Blood samples were collected from dogs belonging to the surveyed households by teams of trained veterinarians. All dogs living in the area since birth or adopted before the age of three months were eligible and included when possible. Samples were taken by venipuncture of *Vena cephalica*, into a tube without anti-coagulant, stored and transported at 4 to 8°C in a cool box, centrifuged in the field and with the resulting sera stored at +4°C, then frozen at −80°C within 3 days at most until laboratory analysis. A free deworming treatment was given to each dog sampled. Age, sex, household location, daily routine as well as dog feeding habits were collected using a structured questionnaire.

### Cattle data

A cross-sectional survey was carried out in cattle in March 2024. Based on an expected prevalence of 19.4% [11], the number of animals to be sampled was 15 cattle per year of birth (for a relative precision of 100% and assuming perfect sensitivity and specificity of the serological test). Due to the rainy season, Tsaratanana commune was inaccessible and excluded from livestock sampling. The remaining four communes were surveyed, focusing on areas where dogs had been sampled in 2023. Only cattle born in the surveyed herd or reported as originating from the survey area were considered for analysis in this study. Farmers were encouraged to present their herd at sampling points designated by local veterinarians for each commune. Some additional herds were identified by snowball sampling. Blood samples were collected by qualified veterinarians by venipuncture of the jugular vein, into a tube without anti-coagulant. Samples were transported at 4 to 8°C, centrifuged and sera stored at - 80°C. All available cattle were sampled with the owner’s consent and deworming agents were given to each owner. Age and sex of each sampled animals were collected.

### Human data

Human dry blood spot (DBS) samples from the 2021 collection of the Ifanadiana Health Outcomes and Prosperity longitudinal Evaluation (IHOPE) cohort were used for this study [38]. The IHOPE cohort was established in 2014 to obtain demographic, health and socio-economic information from a representative sample of 1,600 households in the Ifanadiana district. The Madagascar National Institute of Statistics (INSTAT) was responsible for data collection, survey coordination, training and oversight. Eighty clusters were randomly selected from mapped areas during the 2009 census [39], and twenty households were randomly selected from each cluster. The 2021 wave of data collection (April 22 to June 20) included, for consenting individuals of all ages, a DBS obtained by finger prick using a single-use lancet needle by trained nurses, with 1 to 5 DBS collected on Whatman 903 Protein Saver Card (WHA10531018, SIGMA ALDRICH, EU) filter papers for each individual. All participants (≥ 15 years) provided oral informed consent for the in-person interview and written informed consent for biological sample collection. Parents or guardians provided written consent for biological sample collection from children ≤ 15 years of age, and children 7-14 years provided written consent separately. In our study, only data from individuals living within a radius of less than 5 km from the nearest dog sampling location were included. All data used in this study were de-identified to protect participant confidentiality. There was no target sample size for human samples, as final sample size depended on the available human data which fulfilled these conditions.

### Serological analyses

#### Dogs and cattle sera

Dogs and cattle sera were tested using a commercially available cELISA kit (ID Screen® Rift Valley Fever Competition Multi-species, IDvet Grabels, France). The individual serological status was defined in accordance with the manufacturer’s protocol. In brief, the competition percentage (S/N%) for each sample was calculated using the following formula (Eq.1):

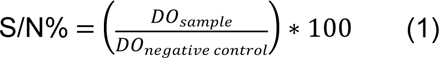

A sample was considered positive if S/N% ≤ 40%, negative if S/N% > 50%, and doubtful if S/N% was between these two values. Doubtful was considered as negative in this study.

#### Human Dry Blood Spots

Antibody detection against the RVFV Nucleoprotein (NP) was performed using Luminex xMAP technology, a bead-based immunoassay (MBA, Luminex®). Human DBS were processed by cutting 3 mm punches, which were eluted overnight at 4°C in phosphate buffered saline (PBS – E404, VWR, USA) - Tween 0.5% (P1379, SIGMA, USA) under gentle shaking. The eluates were then transferred to 1.5 mL tubes and stored at −20°C. DBS eluates were dispensed into designated wells and incubated with MAGPLEX COOH-microsphere beads (MC10012-21, Luminex, USA) coated with NP for 45 minutes. The beads were subsequently placed on a magnetic plate for 60 seconds and washed with an assay buffer (PBS containing 0.05% BSA, 0.1% Tween, pH 7.4). F(ab’)2-Goat anti-Human IgG Fc Secondary Antibody, PE (H10104, Thermo Fisher Scientific, Massachusetts, USA) was added to form antigen-antibody-conjugate complexes. Following a second wash to remove unbound conjugate, fluorescence intensity was measured using a Magpix instrument (MAGPX12234702, Texas, USA). The fluorescence signal, expressed as median fluorescence intensity (MFI), was quantified and was proportional to the amount of specific IgGs in the sample. To account for variability in blood spot volumes across samples, MFI was normalized to total protein content, determined using a Bradford protein assay [40].

In the absence of a well-characterized set of negative and/or positive reference samples, a modelling approach was employed to classify samples as belonging to positive or negative serological status. A finite mixture model was used to identify clusters within the normalized MFI results across the analyzed population [41]. This approach allowed for the classification of individuals as either low or highly reactive. The model outputs included the number of clusters and the distribution of individuals per cluster, based on the probability of belonging to a cluster. A hierarchical clustering model was then applied to consolidate the identified clusters into two primary groups, that were defined as seropositive (highly reactive) and seronegative (low reactive).

MBA allowed to detect anti-RVFV immunoglobulins of type G (IgG) antibodies directed against the RVFV NP, while cELISA detected IgG and IgM antibody responses without differentiation. We assumed a perfect specificity (100%) of these tests in each species [42–44]. All serological assays were performed at the *Institut Pasteur de Madagascar* (IPM).

### Serocatalytic models

Serocatalytic models were implemented to reconstruct the RVFV force of infection (FOI) exerted on individuals of the three studied species, defined as the rate of a susceptible individual becoming infected over time [45]. Assuming that the RVFV antibody response is life-long in dogs, cattle and humans of our sample (*i.e.* children aged less than 10 years) [46–49], the FOI *λ* can be estimated by fitting simple serocatalytic models to serological data of individuals of known age. According to this modelling framework, the serological status *Sero_s,b_* of an individual of species *s* with known birth year *b* follows a Bernoulli distribution (Eq.2):

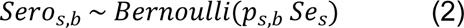

where *Se_s_* is the sensitivity of the serological test used in species *s*, and *p_s,b_* the probability of having been exposed to RVFV prior to time t_s_, the year of sampling in species *s*.

We neglected the probability of seroreversion, and p*_s,b_* was expressed as (Eq.3) [50]:

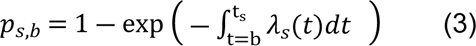

where *λ_s_*(*t*) is the force of infection exerted on individual *s* at time (year) *t*.

#### Hypotheses

Assuming no spatial variation of RVFV FOI within the study area, we tested three hypotheses regarding the relationship between RVFV FOI exerted on dogs (*s = d*) on each year t, *λ*_*d*_(*t*), and on each of the two other species s, *λ*_*s*_(*t*). The null hypothesis (H0) assumed no relationship between *λ*_*d*_(*t*) and *λ*_*s*_(*t*). The first alternative hypothesis (H1) assumed that the FOI exerted on a given species *s* (cattle or humans) was proportional to dog’s FOI (Eq. 4):

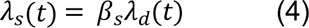

where *β*_*s*_ was the proportionality parameter for species *s*. In the second alternative hypothesis (H2), we took into account the fact that dogs may be contaminated either by mosquito bites or by consuming infectious abortion products from cattle (fetus, placenta). Humans can also be infected through direct contact with tissues and fluids of infected animals (especially slaughterhouse staff and butchers as well as veterinary field officers). However, given that we only used human samples from children under the age of 10 (born after 2010 and sampled in 2021), human exposure was assumed to be vector-borne only. In cattle, although direct transmission is possible, vectorial transmission remains the most important route described in the literature. Therefore, in H2 we assumed that dogs FOI was composed of a vectorial fraction *λ*_*d.v*_(*t*) and a direct transmission fraction *λ*_*d.c*_(*t*), in order to account for the two transmission routes of RVFV. Moreover, the FOI of a given species *s* (cattle or humans) was assumed to be proportional to the vectorial fraction of dog’s FOI (Eq. 5):

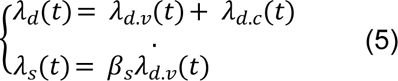

To control for between-years FOI variations, time was divided into 9 intervals: first, because of the low number of individuals exposed during these years, the period from 2011 to 2016 (with FOI assumed constant across these years), and then yearly periods from 2017 to 2024. As the FOI exerted in cattle in 2024 cannot be calculated from the FOI exerted in dogs in H1 and H2 (dog sample collection ended in 2023 while bovine sample collection extended until 2024), it was estimated independently.

The sensitivity of the serological tests and the FOI were hardly identifiable in our models (since sensitivity and exposure probability appeared multiplicatively in equation 2). To allow parameter estimation, we thus made the simplifying assumption of a perfect sensitivity in each species.

#### Model fitting and comparison

Combining the three hypotheses (H0, H1 and H2) for the dog-human and dog-cattle FOI relationships resulted in nine serocatalytic models. According to Eq.2, a binomial likelihood function was formulated to link each model to the data and to estimate the model parameters. Posterior distributions of the parameters were obtained using Bayesian Markov Chain Monte Carlo (MCMC) sampling with RStan plugin (Version 2.32.6) implemented in R (Version 4.4.1). Assuming flat priors for all parameters, 11,000 iterations were performed with 1,000 for initial burn-in iterations and a thinning of 5, using 4 MCMC chains, to obtain the median and 95% credible intervals (95% CrI) of the parameters. The convergence and efficiency of the sampling process were assessed using the diagnostic metrics Rhat (Potential scale reduction factor) and NEFF (Effective sample size) ratio (S2 Table). Autocorrelation was checked graphically with autocorrelation function (ACF, S1 Fig.). Models were compared using the deviance information criterion (DIC), a measure of model fit. The best model was the one with the lowest DIC. The most parsimonious model (with the lowest number of parameters) was preferred when models had similar DIC values (*i.e.* when the DIC difference was below 10) [51].

## Results

Overall, 1,134 individual samples were considered for the study, including 513 dogs, 486 humans and 135 cattle samples. Sampling locations are shown in Fig 1, with 113 villages for dogs, 28 household clusters for humans and 14 sampling points for cattle.

### Seroprevalence

The overall RVFV seropositivity rates were 4.5% (95% CI [2.9 - 6.7]), 17.7% (95% CI [14.4 – 21.4]) and 24.4% (95% CI [17.5 - 32.6]) in dogs, humans and cattle respectively. Maximum seroprevalence of RVFV by species was observed in animals born in 2019 for dogs, 2018 for cattle, and in children born in 2021 (although only 8 children sera were available for this birth year). Serological results by year of birth are presented in Table 1.

**Table 1.**
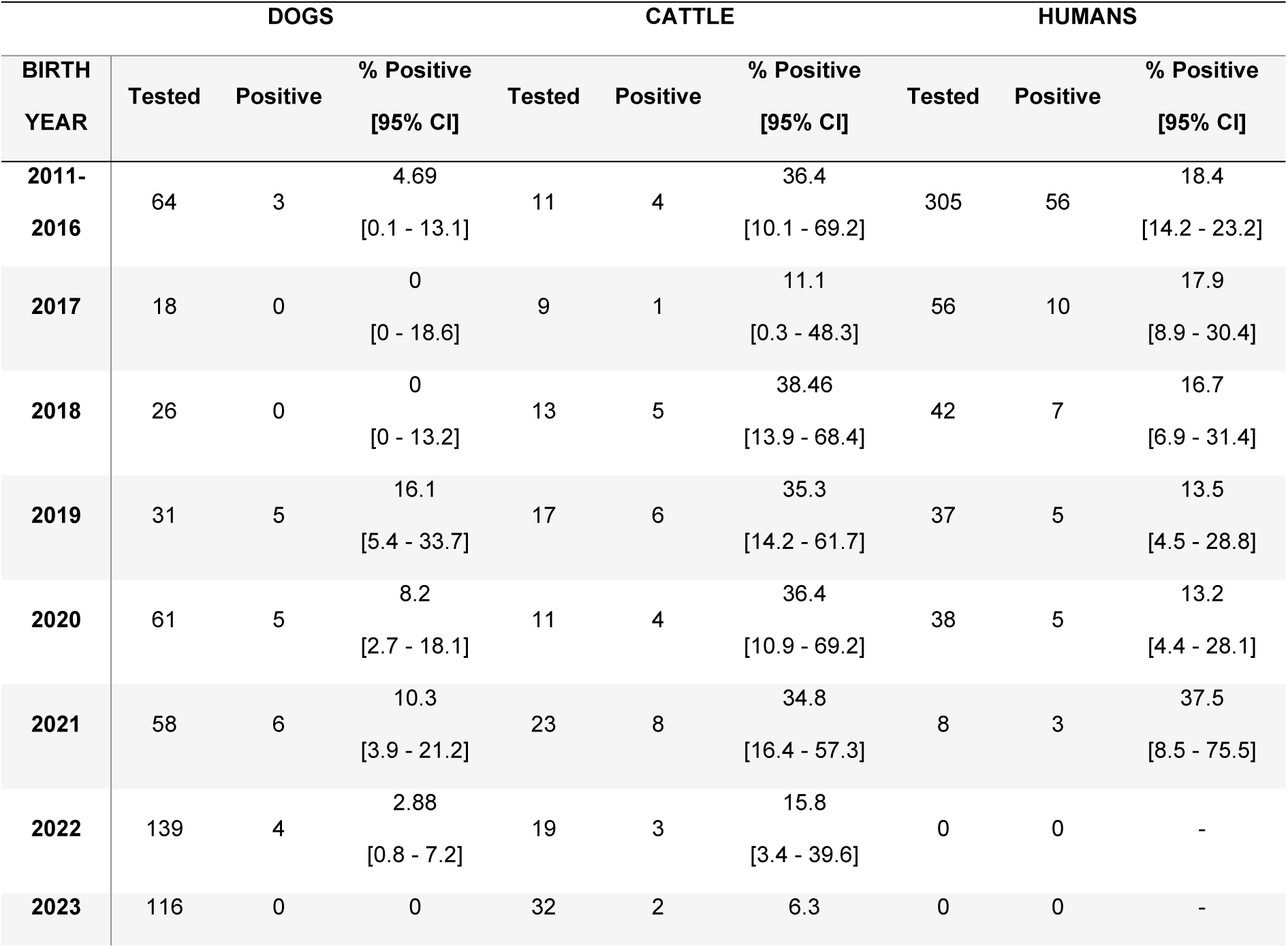
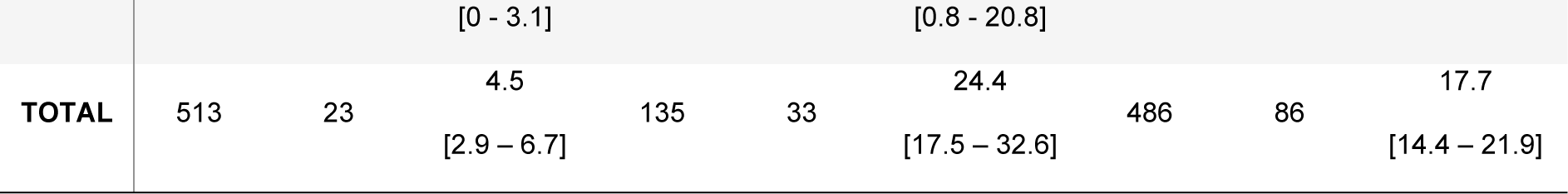
Number of individuals tested and serological results by year of birth (or category for individuals born between 2011 and 2016) and by species.

For dogs, the seroprevalence by commune varied between 2.1% (95% CI: [0.4 – 5.8], Ifanadiana) and 10.9% (95% CI: [4.5 – 21.6], Tsaratanana). For cattle, it varied between 16.2% (95% CI: [6.2 - 32.0], Kelilalina) and 33.3% (95% CI: [18.0 - 51.8], Antaretra). For humans, it varied between 9.2% (95% CI: [4.1 - 17.3], Kelilalina) and 16.1% (95% CI: [9.3 - 25.2], Antaretra). No consistent spatial seroprevalence trend was visually observed in the three species (Fig. 2, S3 Table).

**Fig. 2.**
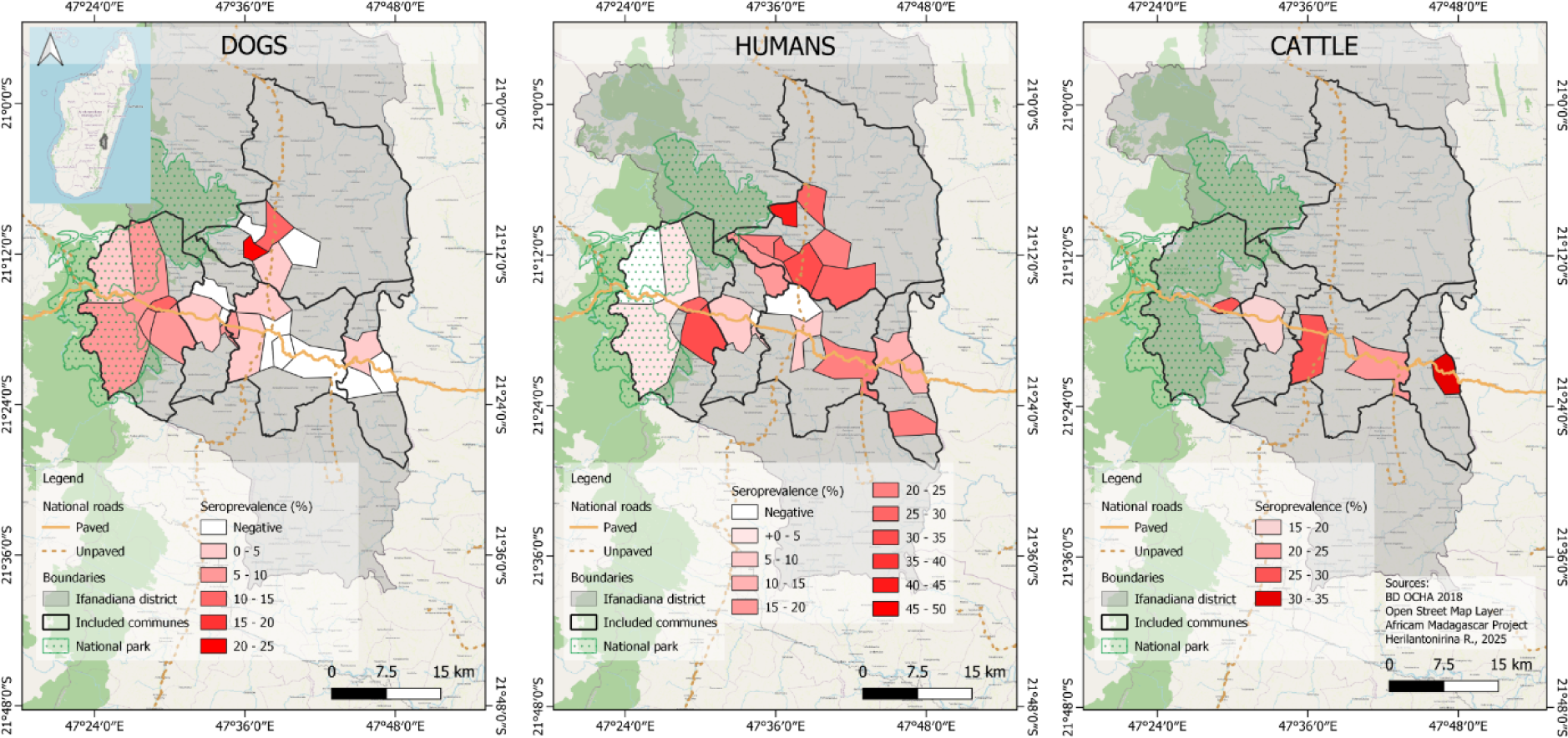
Spatial distribution of Rift Valley fever virus seroprevalence in dogs (2023), humans (2021) and cattle (2024) in the Ifanadiana district, Madagascar.

### Serocatalytic models

We fitted 9 serocatalytic models to the serological data, for which DIC values and number of estimated parameters are provided in Table 2.

**Table 2.**
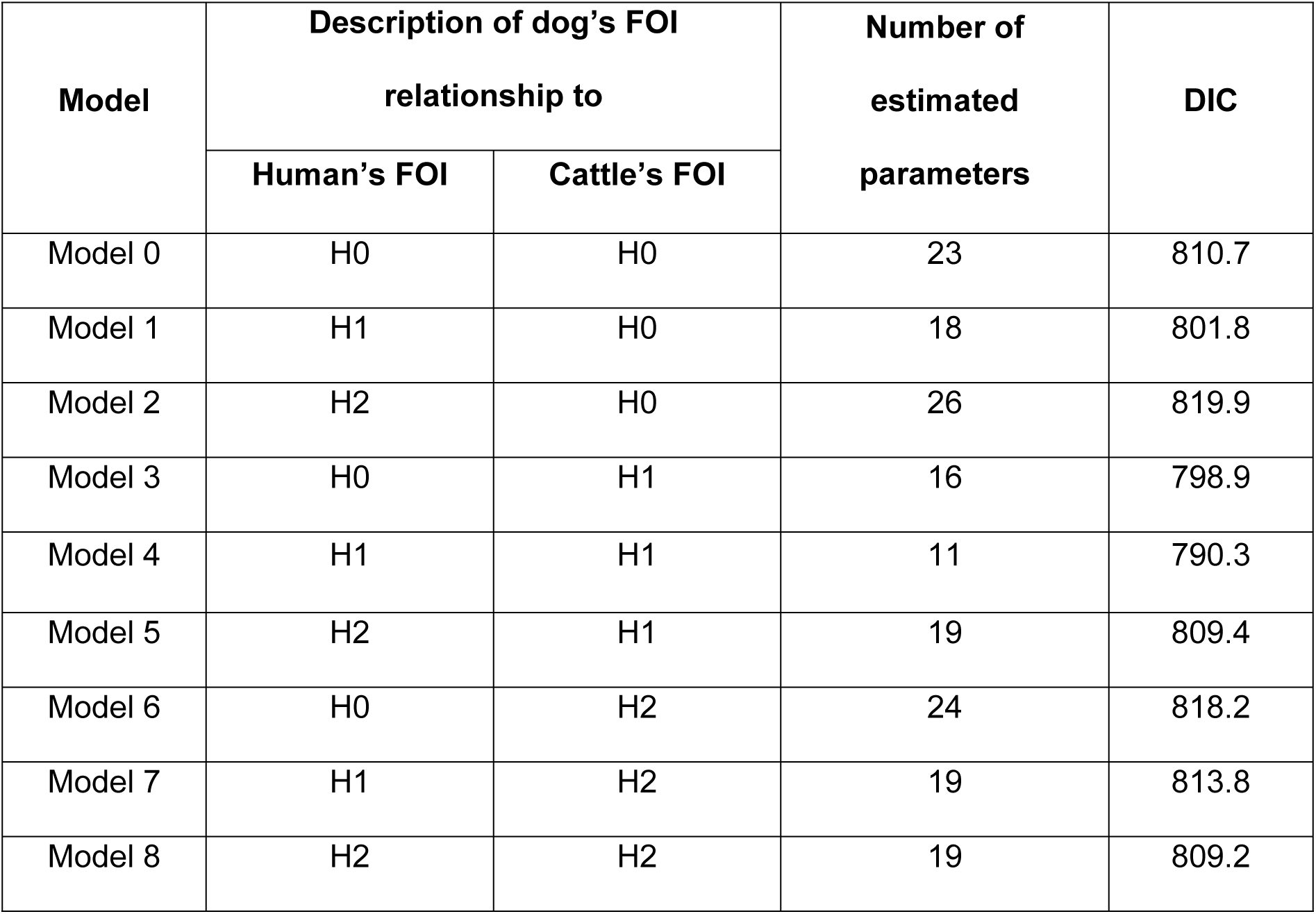
Description of serocatalytic models, with corresponding number of estimated parameters and deviance information criterion (DIC) values. The human force of infection (FOI) (respectively the cattle FOI) is modelled as proportional to the dog’s FOI under hypothesis H1, to its vector-borne fraction only under hypothesis H2, or unrelated under null hypothesis H0.

| Model | Description of dog's FOI relationship to |  | Number of estimated parameters | DIC |
| --- | --- | --- | --- | --- |
|  | Human's FOI | Cattle's FOI |  |  |
| Model 0 | H0 | H0 | 23 | 810.7 |
| Model 1 | H1 | H0 | 18 | 801.8 |
| Model 2 | H2 | H0 | 26 | 819.9 |
| Model 3 | H0 | H1 | 16 | 798.9 |
| Model 4 | H1 | H1 | 11 | 790.3 |
| Model 5 | H2 | H1 | 19 | 809.4 |
| Model 6 | H0 | H2 | 24 | 818.2 |
| Model 7 | H1 | H2 | 19 | 813.8 |
| Model 8 | H2 | H2 | 19 | 809.2 |

Models 3 and 4 had the lowest DIC values (with less than 10 points difference between them). Since model 4 was more parsimonious than model 3 (11 estimated parameters for model 4 against 16 for model 3), we selected model 4, in which human’s and cattle’s FOIs were both proportional to dogs’ FOI. In dog, before 2021, the estimated median FOI ranged between 0.003 (95% CrI: [7 10^-5^ – 0.009]) and 0.009 (95% CrI: [2.5 10^-4^ – 0.033]). We observed a significant increase in dog’s FOI in 2021, with a median value of 0.048 (95% CrI: [0.018 – 0.086]), followed by a decrease in 2022 (0.023, 95% CrI: [0.004 – 0.049]) and 2023 (0.006, 95% CrI: [1.8 10^-4^ – 0.021]). By construction, FOIs calculated for humans (until 2021, year of sampling) and cattle (until 2023) followed the same trends as dog FOI, with *β_h_* and *β_b_* proportionality parameters estimated at 2.6 (95% CrI: [1.4 – 5.1]) and 3.5 (95% CrI: [1.2 – 6.4]) respectively (Fig 3).

**Fig 3.**
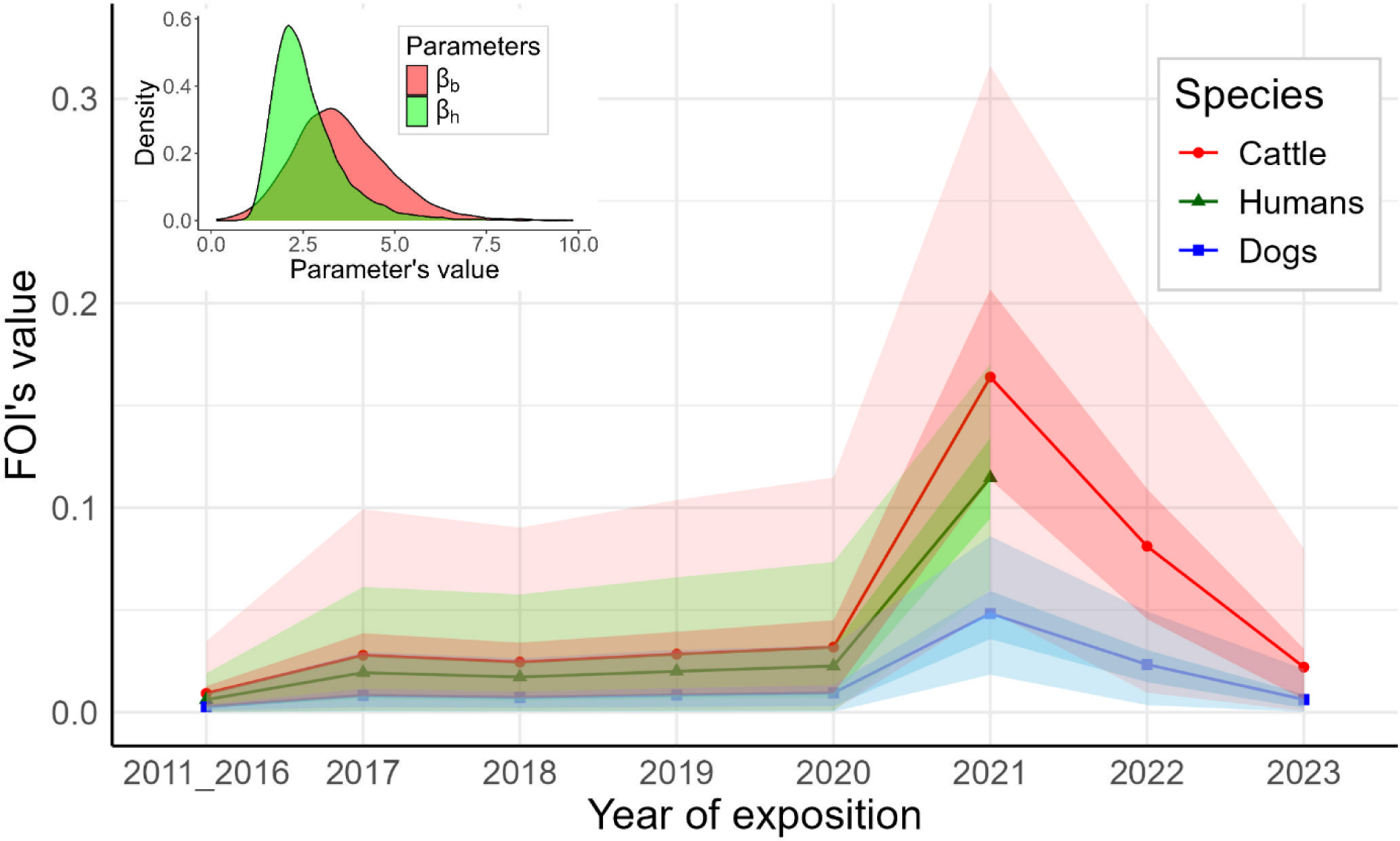
Estimates of the annual RVF FOI in dogs, humans, and cattle according to the best fitting serocatalytic model (Model 4), with FOI in cattle and human proportional to dog’s FOI), Ifanadiana district, Madagascar

The predicted values of RVFV seroprevalence fitted adequately the observed data: whatever the year of birth, the observed seroprevalence was within the credible intervals of the predicted value (Fig 4).

**Fig 4.**
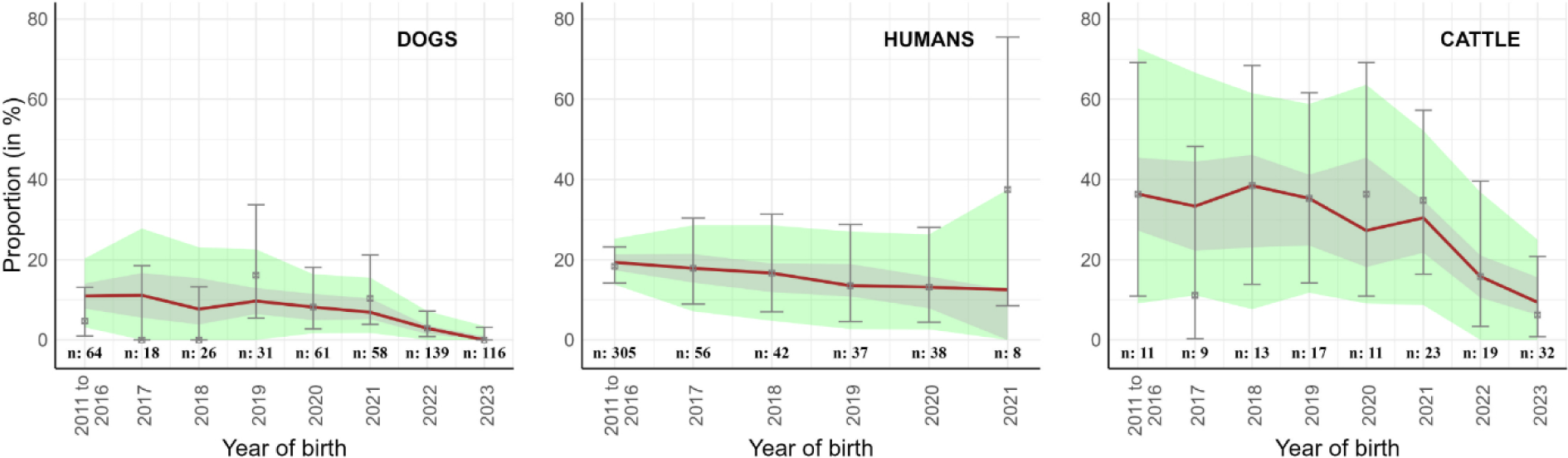
Predicted versus observed seroprevalence of RVFV in dogs, humans and cattle in Ifanadiana district. Grey and green colors represent the 50% and 95% bayesian credible intervals of predicted seroprevalence values, respectively, while red lines represent the median values. Points represent the observed seroprevalence and error bars the corresponding 95% confidence interval. The number of samples per year of birth is indicated by “n” values

## Discussion

In our study, we estimated the RVFV FOI in dogs, humans, and cattle, using serocatalytic models applied to age-structured serological data [50], and analyzed the quantitative relationship between dog’s FOI and the FOIs exerted on cattle and humans over time. The model that most accurately described the observed serological results (on the basis of the DIC) linked canine FOI to bovine and human FOIs by two linear relationships, with proportionality coefficients estimated at 3.5 and 2.6, respectively. This model identified a RVFV FOI peak in 2021, which is the year of the last RVF outbreak in the Mananjary district, the neighboring district of our study site in the south-east of Madagascar. This finding suggests that there was a significant RVFV circulation during this outbreak in the Ifanadiana district, although no clinical cases were reported either in cattle nor humans. To our knowledge, this study reports the first serological detection of RVFV in dogs in Madagascar, and is the first attempt to estimate the FOI of RVFV in dogs and its quantitative relationship with RVFV FOIs in humans and other exposed species.

The seroprevalence we observed in Malagasy dogs was comparable to the 3.8% (1/26) observed in dogs sampled in 2018-19 in a forest area in Gabon [28]. In Egypt, one out of four sampled dogs were reported as seropositive, during the 1977-78 RVF outbreak [29]. Among the three species here studied, the highest seroprevalence was observed in cattle. Similar differences of seroprevalence between cattle and humans were reported in Madagascar during the 2021 RVF outbreak [2], and during the inter-epidemic/epizootic period after the 2008 outbreak [52,53]. These differences between species are consistent with the RVFV epidemiology [54], given that ruminants are considered to be the main reservoir and amplifying hosts of the virus [55,56], while other species such as humans and dogs are considered as accidental and dead-end hosts [29,57]. Moreover, these differences may be due to several factors including body surface area, attractivity and feeding behaviors of mosquitoes in the area of concern. Indeed, cattle present a larger body mass and surface area, which increases the amount of CO_2_ emitted and the attractiveness of mosquitoes compared to human and dogs. While humans are preferred to dogs by some mosquito species such as *Aedes albopictus* and *Ae. aegypti* [58,59], the opposite is true for other mosquito species such as *Culex quinquefasciatus* [60].

We assumed 100% specificity in our serological tests. Indeed, according to a ring trial performed by Kortekaas *et al.*, the cELISA kit (IDVet), which appears to be widely used for RVFV surveillance programs and among the recommended kit for RVFV serological diagnosis [42], demonstrated high specificity in ruminants (100%) [61]. In addition, to our knowledge, the only phlebovirus whose circulation has been reported in Madagascar is RVFV. Our hypothesis of 100% sensitivity of serological tests was a simplifying assumption made to allow parameter estimation. It was supported by the fact that the cELISA kit (IDVet), which is a multi-species one, has shown high sensitivity in ruminants (ranging from 91 to 100% with mean value of 97.2%) according to Kortekaas *et al.* [61]. Lower sensitivities of the serological tests in humans and/or dogs would have had an impact on the estimates of the proportionality coefficients *β*_*b*_ and *β_h_*, particularly if the sensitivity ratio dog/man (for *β_h_*) or dog/cattle (for *β*_*b*_) departed markedly from 1. Nevertheless, it would not have impacted the model selection procedure. From the 9 tested serocatalytic models, the one that provided the best fit assumed that FOIs in human and in cattle were both proportional to the FOI in dogs. Estimated FOIs in these three species showed very little variation before 2021, suggesting that the virus circulated at a low level during these years (Fig 3). These findings corroborate previous results, showing a very low-level transmission during inter-epidemic/inter-epizootic periods in Madagascar [11,13–15]. Olive *et al* have shown that RVFV FOI in Malagasy ruminants varied over time but also by ecological regions [62]. As all municipalities in our study were from the same district (Ifanadiana), we assumed that there was no spatial variation in RVFV FOIs at the district scale. Although the abundance and diversity of mosquitoes may vary from the rainforest to rainforest edge and savanna biotope, which could result in significant variation of RVFV vectorial transmission [63], there was no consistent spatial trend in RVFV seroprevalence across host species in our study (Fig 2).

We chose to build models with a time-varying FOI to control for the yearly variations of FOIs. Given the limited number of samples from cattle and dogs born between 2011 and 2016, and to address the potential bias of precision in the year of birth of these older animals, we assumed that the FOIs through these years were constant. Keeping these data allowed us to have more statistical power and precision in parameter estimations [62]. Despite this assumption, our estimate of RVFV FOIs exerted on cattle from 2011 to 2016 (0.009; 95% CrI: [2.10^-4^ – 0.035]) was within the median range of inter-epizootic RVFV FOIs estimated values from mid-2010 to mid-2014 (0.009 to 0.033) in the Eastern eco-regions of Madagascar, described in cattle in a previous study [62]. Given that RVFV infection confers long-lasting immunity of more than 20 years in humans, and is considered lifelong in ruminants [46,64], we assumed that all exposed individuals that were infected (including dogs) would remain positive for life without seroreversion, or that at least the RVF antibody response would not wane until the age at sampling (maximum 12 years), taking into account a continual exposure to the RVFV in an endemic area as Madagascar.

The use of dogs for serological surveillance of zoonotic arboviruses in Madagascar has the potential to be cost-effective [33], as a single population sample could be used to detect the presence and the circulation of multiple pathogens, using several specific commercial or in-house serological tests, including multiplex techniques. Indeed, besides RVFV, the circulation of several arboviruses, including West Nile virus, Usutu virus, Dengue virus, and Chikungunya virus, has been documented in the island [65,66]. Moreover, the potential use of dogs as sentinel animals for up to 53 pathogens was described in the literature, and it was already assessed and suggested for *Yersinia pestis* in Madagascar [23,67].

In conclusion, the existence of a quantitative relationship between dogs, humans and cattle FOIs demonstrated here suggests that sampling dogs could allow to identify and prioritize risk areas for RVF, when human serological data are not available, and access to ruminants is limited. However, further studies in other ecological regions are needed to confirm the existence of this proportional relationship between dog, human and cattle FOIs, and to document the variations of the proportionality parameters, according to the vector diversity in these regions, their relative abundance, and trophic preferences, but also according to dogs’ lifestyles.

## Acknowledgments

This work was supported by AfriCAM project, funded by the French Development Agency (AFD) as part of PREACTS program. Data collection for the IHOPE cohort was supported by internal funding from Pivot and support from the Herrnstein Family Foundation. Dog and cattle serosurveys were piloted by the DRZVP/FOFIFA, with CIRAD, IPM, DSV and PIVOT/IRD. We especially thank the dog owners and cattle herders of the Ifanadiana district who voluntarily accepted to participate in the study. Many thanks also to administrative officers (*Fokontany* chiefs and Mayor of communes), veterinarians and community health workers who greatly facilitated our fieldwork. We are grateful to Ann Miller who shared with us the necessary human’s metadata according to the Pivot data sharing policy. At last, but not least, we thank all the technicians for their invaluable assistance with animal sampling and serological analyses including Jean Pierre Ravalohery and Rondroharivelo Lucia Rasoahanitralisoa (*Unité de Virologie – IPM*).

## Data availability statement

Data is presented within the paper and its supplementary material.

## Author contributions

**Conceptualization:** Veronique Chevalier, Benoit Durand, Herilantonirina Solotiana Ramaroson.

**Data curation:** Herilantonirina Solotiana Ramaroson, Solohery Lalaina Razafimahatratra, Laure Chevalier.

**Formal analysis:** Herilantonirina Solotiana Ramaroson, Benoit Durand.

**Funding acquisition:** Veronique Chevalier, Andres Garchitorena.

**Investigation:** Herilantonirina Solotiana Ramaroson, Katerina Albrechtova, Laure Chevalier, Domoina Rakotomanana, Patrick de Valois Rasamoel, Heritiana Fanomezantsoa Andriamahefa, Mamitiana Aimé Andriamananjara.

**Methodology:** Herilantonirina Solotiana Ramaroson, Veronique Chevalier, Benoit Durand, Jonathan Bastard, Vincent Lacoste, Soa Fy Andriamandimby, Matthieu Schoenhals, Andres Garchitorena, Lova Tsikiniaina Rasoloharimanana, Solohery Lalaina Razafimahatratra.

**Project Administration:** Veronique Chevalier

**Resources:** Veronique Chevalier, Benoit Durand, Matthieu Schoenhals, Andres Garchitorena, Lova Tsikiniaina Rasoloharimanana, Solohery Lalaina Razafimahatratra, Katerina Albrechtova, Laure Chevalier, Herilantonirina Solotiana Ramaroson.

**Supervision:** Veronique Chevalier, Benoit Durand, Modestine Raliniaina, Claude Arsène Ratsimbasoa.

**Validation:** Benoit Durand, Veronique Chevalier.

**Visulaization:** Herilantonirina Solotiana Ramaroson.

**Wrtiting – Original Draft Preparation:** Herilantonirina Solotiana Ramaroson, Veronique Chevalier, Benoit Durand.

**Writing – Review & Editing:** all co-authors cited in this article.

## Supporting information Captions

**S1 Table. Dog, human and cattle database with age and year of exposure to Rift Valley fever virus.** (XLSX).

**S2 Table. Effective sample size (NEEF), NEEF ratio, Rhat, and values of estimated parameters per model.** Estimated parameter values, NEEF and Rhat values are presented. Details of parameters: **FOI** is the force of infection, **B** is bovine (cattle), **C** is canine (dogs), **H** is human, **_16** is the 2011 to 2016 years, **17 to 24** are the years 2017 to 2024, **CC** is the contact fraction of dog’s FOI, **CV** is the vectorial fraction of dog’s FOI. **BBETA** and **HBETA** are the estimated proportionality parameters for bovine and human FOIs, respectively, to that of dogs. (XLSX).

**S3 Table**. **Individuals tested and positive for anti-RVFV antibodies by communes and species, with Chi-2 / Fisher exact p-values.** In dogs and humans, seroprevalence in Tsaratanana was the highest, with 10.9% (7/64, 95% CI: [4.5 – 21.2]) and 29.5% (43/146, 95% CI: [22.2 – 37.6]), respectively. However, without considering Tsaratanana, no spatial difference in seroprevalence was found between localities in the three species. (XLSX).

**S1 Fig. An example of autocorrelation graph from autocorrelation function (ACF), model 4**. Autocorrelation graph for the selected model 4 (rows: the four MCMC chains with lag = 10, columns: estimated parameters). (PDF).

## Notes

### Competing Interest Statement

The authors have declared no competing interest.

